# LGBTQ+ realities in the biological sciences

**DOI:** 10.1101/2025.01.24.634486

**Authors:** Katelyn M. Cooper, Carly A. Busch, Alice Accorsi, Derek A. Applewhite, Parth B. Bhanderi, Bruno da Rocha-Azevedo, Abhijit Deb Roy, Joseph P. Campanale, Fred Chang, Jerry E. Chipuk, Lee A. Ligon, G.W. Gant Luxton, Austin J. Graham, Camila Hochman-Mendez, Imge Ozugergin, Zachory M. Park, Claire M. Thomas, Alex M. Valm, Hongxian Zhu, Rebecca S. Alvania

**Affiliations:** LGBTQ+ Committee, The American Society for Cell Biology; Rockville, MD 20852, USA; Research for Inclusive STEM Education Center, School of Life Sciences, Arizona State University; Tempe, AZ 85281, USA; Department of Molecular and Cellular Biology, College of Biological Sciences, University of California Davis; Davis, CA 95616, USA; Department of Biology, Reed College; Portland, OR 97202, USA; Research Operations, Eikon Therapeutics; Hayward, CA 94545 USA; Center for Cell Analysis and Modeling, Department of Cell Biology, University of Connecticut School of Medicine; Farmington, CT – 06030, USA; School of Life Sciences, University of Nevada Las Vegas, Las Vegas, NV 89154 USA; Department of Cell and Tissue Biology, University of California San Francisco; San Francisco, CA 94143 USA; The Tisch Cancer Institute at the Icahn School of Medicine at Mount Sinai, New York City, New York, 10029 USA; Department of Biological Sciences and Center for Biotechnology and Interdisciplinary Studies, Rensselaer Polytechnic Institute; Troy, NY 12180 USA; Department of Pharmaceutical Chemistry, University of California, San Francisco; San Francisco, CA 94158, USA; Regenerative Medicine Research, The Texas Heart Institute; Houston, TX 77030, USA; Institut Pasteur; Paris, 75015, France; Department of Biology, Georgetown University; Washington, DC 20057, USA; Department of Biology and Department of Biochemistry and Molecular Biology, The Pennsylvania State University; University Park, PA 16802, USA; Department of Biological Sciences and RNA Institute, University at Albany State University of New York; Albany, NY 12222, USA; Department of Molecular Genetics, University of Toronto; Toronto, M5S 1A8, Canada; Executive Office, The American Society for Cell Biology; Rockville, MD 20852, USA

## Abstract

While scientific environments have been described as unwelcoming to the LGBTQ+ community, and fields such as physics have systematically documented these challenges, the climate in biology workplaces has not been assessed. We conducted the largest survey to date of LGBTQ+ biologists to examine how their sense of belonging and perception of climate in the biology workplace and professional societies compare to that of their straight and cis peers. We surveyed 1419 biologists across five professional societies, with 486 identifying as LGBTQ+. Trans and gender non-conforming (TGNC) biologists reported lower belonging and morale within the workplace, professional societies, and the biology community compared to cis, straight biologists. They also reported being less comfortable with the climate of various professional biology environments. While LGBTQ+ biologists report that their workplaces are moderately inclusive, over 20% of all LGBTQ+ biologists and nearly 40% of TGNC biologists experience exclusionary behavior at work. This landmark survey provides the first comprehensive analysis of the LGBTQ+ climate in biology, revealing specific challenges faced by TGNC scientists and identifying interventions to enhance inclusivity for scientists.

**Significance Statement:** This landmark study includes the largest known sample of LGBTQ+ biologists and offers the first comprehensive description of the LGBTQ+ climate in biology, differentiating between the experiences of cisgender lesbian, gay, bisexual, and queer (LGBQ) biologists and transgender and gender non-conforming (TGNC) biologists. The study found that compared to non-LGBTQ+ biologists, TGNC participants report lower belonging, morale and comfort with the climate across biology workplaces, professional societies, and the biology community. While on average LGBTQ+ participants reported that their workplaces are moderately inclusive, over 20% of all LGBTQ+ biologists and nearly 40% of TGNC biologists report experiencing exclusionary behaviors at work. The study offers immediate implications for institutional policies and professional development in the biological sciences.

## Introduction

Individuals who identify as lesbian, gay, bisexual, transgender, queer/questioning plus (LGBTQ+) have historically encountered discrimination and hostile environments in science, technology, engineering, and mathematics (STEM) fields (1–3). The unwelcoming nature of the STEM community is attributed to the pervasive cisheteronormativity (4–6) and the cultivation of an apolitical culture that sidelines diversity issues (2, 7, 8). Specifically, LGBTQ+ individuals have previously described a spectrum of negative experiences in STEM environments. For example, physicists have reported experiencing discrimination due to their LGBTQ+ identities (9), LGBTQ+ environmental scientists reported being disrespected by their peers (10), and LGBTQ+ STEM professionals reported being devalued by their colleagues and socially excluded (2). Further, LGBTQ+ scientists are likely underrepresented and underserved in STEM (11–13), yet some scientific organizations do not consider LGBTQ+ individuals to be members of an underrepresented group (14).

There have been some efforts to systematically assess the experiences of LGBTQ+ individuals in professional STEM environments. These studies primarily surveyed individuals across STEM disciplines (15, 16), with the most recent large-scale study in 2021 finding that LGBTQ+ STEM professionals experience career limitations, harassment, and professional devaluation to a greater extent than their non-LGBTQ+ peers (2). Discipline-specific assessments are rare, with the 2016 LGBT+ Climate in Physics Report (17) being a notable exception. This study of 324 LGBT+ physicists concluded that the climate experienced by LGBT+ physicists was highly variable. Social norms often established expectations for physicists to conceal their queer identities and many scientists reported feeling isolated and experienced or observed exclusionary behaviors toward LGBTQ+ individuals in the workplace. Considering the cultural differences among STEM fields and varying efforts to create inclusive environments, there is a pressing need for additional, discipline-specific evaluations of LGBTQ+ climate.

The life sciences provide a particularly important context in which to assess the LGBTQ+ climate, given the potential intersection of biology content and the LGBTQ+ identity. Many biologists regularly encounter pervasive myths regarding the biology of attraction, biology of gender, genetics, reproduction, hormones, and sexuality, and may be expected to teach content that has been historically influenced by binary and cisheteronormative thought (18, 19). Interview studies have found that some LGBTQ+ undergraduates (20) and graduates (21) describe biology classrooms and graduate programs as unwelcoming and unaccepting spaces and some LGBTQ+ biology professors highlight the potential for negative repercussions if they were to reveal their identity within their department (22). The intersection between content and LGBTQ+ identities uniquely positions the biology community to lead changes to make STEM more inclusive. Yet, no large-scale studies have specifically probed the experiences of LGBTQ+ biologists as it relates to their work environments and professional societies.

The experiences of trans and gender non-conforming (TGNC) biologists are particularly important to understand, given that TGNC individuals encounter more extreme harassment, discrimination, and violence than their cis LGBQ counterparts (23–25). In biology, there are unique ways in which the TGNC identity intersects with biology curriculum; namely, gender is often discussed in ways that erase or marginalize non-cisnormative identities (23, 26, 27). An interview study of TGNC biology students revealed that biology can be implicitly exclusionary and negatively affects their sense of belonging, career preparation, and interest in biology content (27). However, there is a paucity of research that considers gender beyond the binary (28), particularly when studying the experiences of biologists (23).

To date, no large-scale study has compared the experiences of cis LGBQ biologists, TGNC biologists, and non-LGBTQ+ biologists, nor documented the climate of biology environments for cis LGBQ and TGNC biologists. To address this gap, we surveyed 1419 biologists across five professional societies, 486 of whom identify as LGBTQ+ (Table S1), capturing identities across the spectrum including 153 TGNC biologists. We aimed to assess: 1. To what extent does the belonging, morale, and comfort of LGBTQ+ and non-LGBTQ+ biologists differ? 2. How inclusive are biology workplaces for LGBTQ+ individuals? 3. To what extent do LGBTQ+ biologists experience discrimination? 4. To what extent are LGBTQ+ individuals out in different biology environments? 5. What recommendations do LGBTQ+ biologists have for creating more inclusive biology environments? When answering our research questions, our large sample size enabled us to answer these research questions with unprecedented depth by distinguishing between the unique experiences of cis LGBQ and TGNC biologists.

## RESULTS

In reporting the results, we include any participant who indicated that they were a part of the LGBTQ+ community as “LGBTQ+.” In analyses, we considered cisgender LGBQ biologists separately from TGNC biologists. The “cis LGBQ” group includes any participant who selected an orientation identity (e.g., lesbian, gay, bisexual) and reported that they are cisgender. The “TGNC” group includes individuals who did not select that they are cisgender, such as participants who selected agender, genderqueer, gender nonbinary, or transgender. Individuals within the TGNC group may also hold orientation identities. This distinction allows us to identify unique challenges faced by different members of the LGBTQ+ community.

### TGNC biologists report lower belonging, morale, and comfort with climate across a variety of professional contexts

Compared to cisgender straight biologists, and controlling for race, professional position, and geographic location, TGNC biologists reported lower sense of belonging and morale across professional contexts including their workplace (belonging: β = -1.00, p < .0001; morale: β = -0.71, p = .004), within the field of biology (belonging: β = -1.05, p < .0001; morale: β = -1.02, p < .0001), and in their professional societies (belonging: β = -1.08, p = .0002; morale: β = -0.93, p = .0002). TGNC individuals also perceived less comfortable climates on campus (β = -0.84, p < .0001; Fig. 1B), in their departments (β = -0.58, p = .002), and as students (β = -1.28, p = .0003) compared to cisgender straight participants.

**Figure 1.**
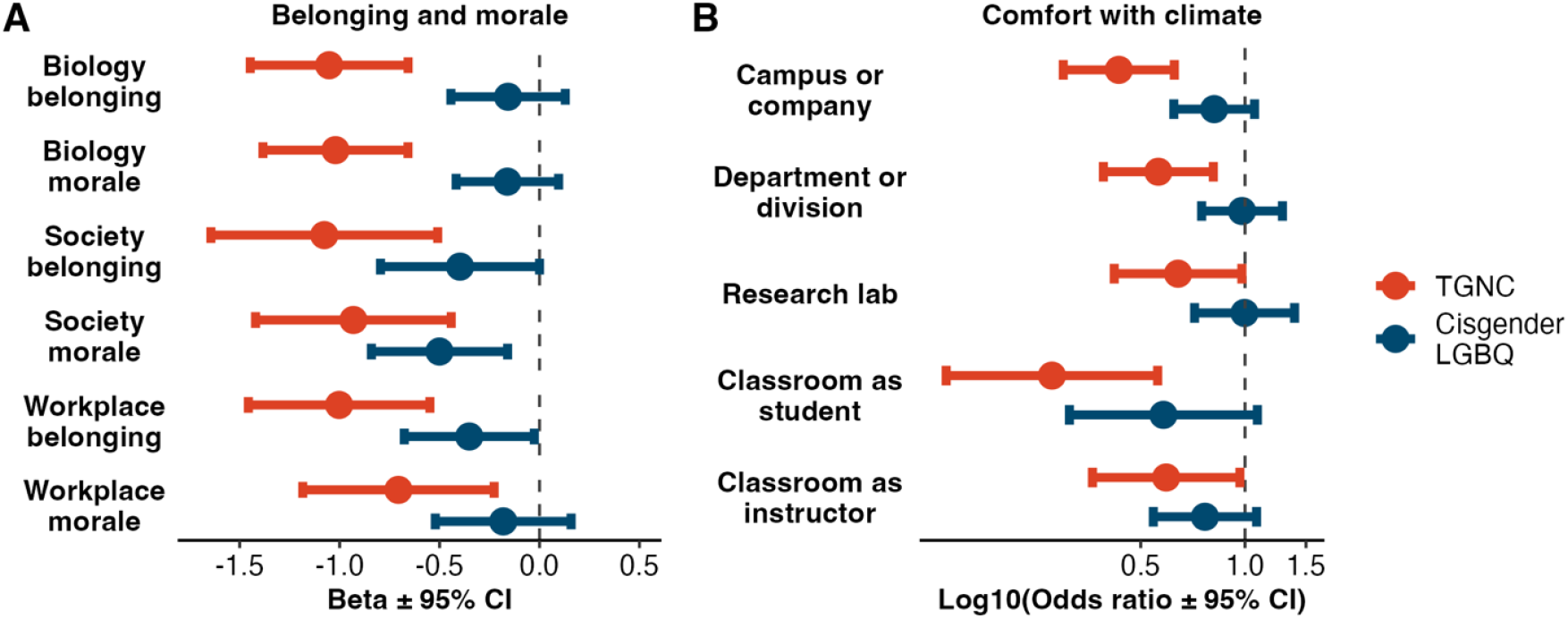
Forest plot of regression results from: (**A**) linear regressions of sense of belonging and feelings of morale in the biology community, professional societies, and workplace; and (**B**) ordinal regressions for comfort with climate across five common contexts. TGNC participants are represented in red and cisgender LGBQ participants in blue. Points and confidence intervals to the left of the vertical dashed line indicate that the represented group expressed a lower score on the respective outcome relative to the reference group (which for all models is cisgender straight biologists). Confidence intervals which do not cross the vertical dashed line are statistically significant.

Cisgender LGBQ biologists reported lower feelings of morale in their professional societies (β = - 0.50, p = .004), compared to cisgender straight biologists, controlling for gender, race, professional position, and geographic location. There were no significant differences in feelings of belonging and morale within the workplace or field of biology and no significant differences in comfort with climate across contexts between cisgender LGBQ and straight cisgender biologists. Full results from both sets of regressions are available in tables S2 and S3.

### Biology workplaces show mixed levels of LGBTQ+ inclusion and exclusion

The LGBTQ+ workplace climate scale consists of two factors: LGBTQ+ inclusion and LGBTQ+ exclusion. Of the individual items that comprise the inclusion factor (listed in Fig. 2), over half the participants agreed with most statements indicating inclusive workplace environments. However, less than half of the participants agreed that “Non-LGBTQ+ employees are comfortable engaging in LGBTQ+ friendly humor with LGBTQ+ employees,” “Employee LGBTQ+ identity does not seem to be an issue,” and “Coworkers are as likely to ask nice, interested questions about LGBTQ+ relationships as they are about non-LGBTQ+ relationships.” The 5-point scale measuring LGBTQ+ climate inclusion ranged from 1, strongly disagree to 5, strongly agree, with higher scores indicating greater perceived inclusivity. The average score across all items for all LGBTQ+ participants was 3.5. Cisgender LGBQ participants had a mean score of 3.6, while TGNC participants’ mean score was 3.4. When controlling for race, professional position, and geographic location, TGNC participants reported significantly lower inclusion compared to cis LGBQ participants (β = -0.21, *p* = .005).

**Figure 2.**
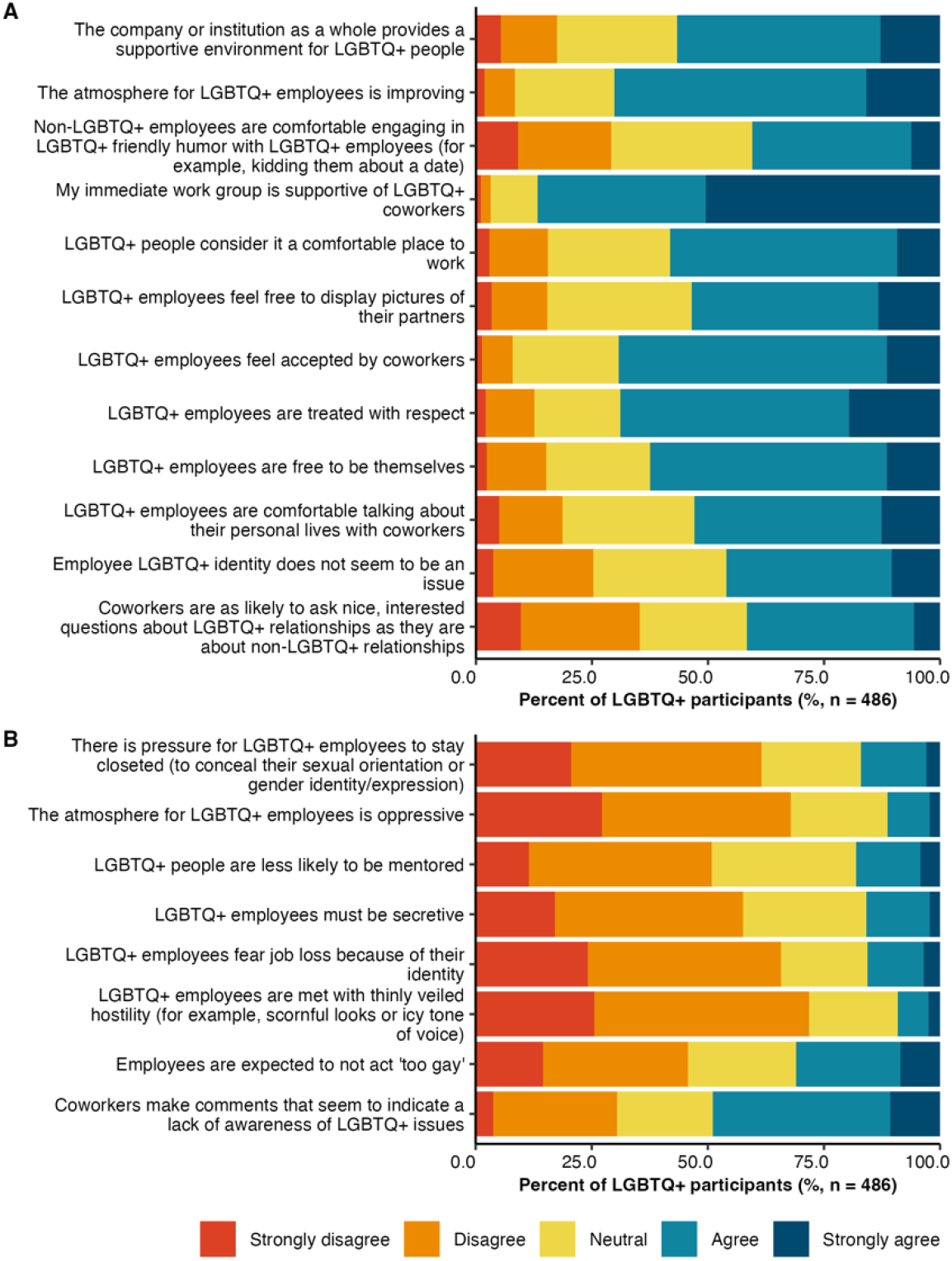
Distribution of responses from LGBTQ+ participants for the: **(A)** 12 positively worded LGBTQ+ inclusion items; and **(B)** 8 negatively worded LGBTQ+ exclusion items assessing department/workplace climate.

Of the individual items that comprise the exclusion factor (listed in Fig. 2), over half of participants disagreed with most statements suggesting relatively inclusive workplace environments. However, less than half of participants disagreed that “Employees are expected to not act ‘too gay’,” and “Coworkers make comments that seem to indicate a lack of awareness of LGBTQ+ issues.” The 5-point scale measuring LGBTQ+ climate inclusion ranged from 1, strongly disagree to 5, strongly agree, where lower scores indicate greater perceived inclusivity. The average score for all items among all LGBTQ+ participants was 2.5 out of 5, Cisgender LGBQ participants averaged 2.4, and TGNC participants reported an average of 2.6. When controlling for race, professional position, and geographic location, TGNC biologists reported higher exclusivity scores than cis LGBQ biologists (β = 0.16, *p* = .06). The distribution of responses for LGBTQ+ inclusion and exclusion climates across all participants is shown in Fig. 2 and full results from both regressions in table S4.

### Over 20% of all LGBTQ+ biologists experience exclusionary behavior at work

The contrast between moderate workplace inclusivity ratings and the actual experiences of LGBTQ+ participants is striking: 21.2% of all LGBTQ+ participants personally experienced intimidating, offensive, and/or hostile conduct due to their LGBTQ+ identity, which interfered with their ability to work or learn in their department or workplace within the past year. Additionally, 38.5% observed or were made aware of such conduct directed toward others. Over forty percent (43.6%) personally experienced exclusionary, intimidating, offensive and/or hostile conduct outside of their department or workplace because of their LGBTQ+ identity within the past year.

When examining the experiences of TGNC biologists specifically, we found that nearly 40% experienced exclusionary behavior at work and over 50% experienced it outside of work (Fig. 3). When controlling for race, professional position, and geographic location, TGNC biologists were significantly more likely to report experiencing exclusionary behavior at work (odds ratio (OR) = 3.73, *p* < .0001) and outside of work (OR = 2.67, *p* < .0001) compared to cis LGBQ biologists. Full regression results are available in table S5.

**Figure 3.**
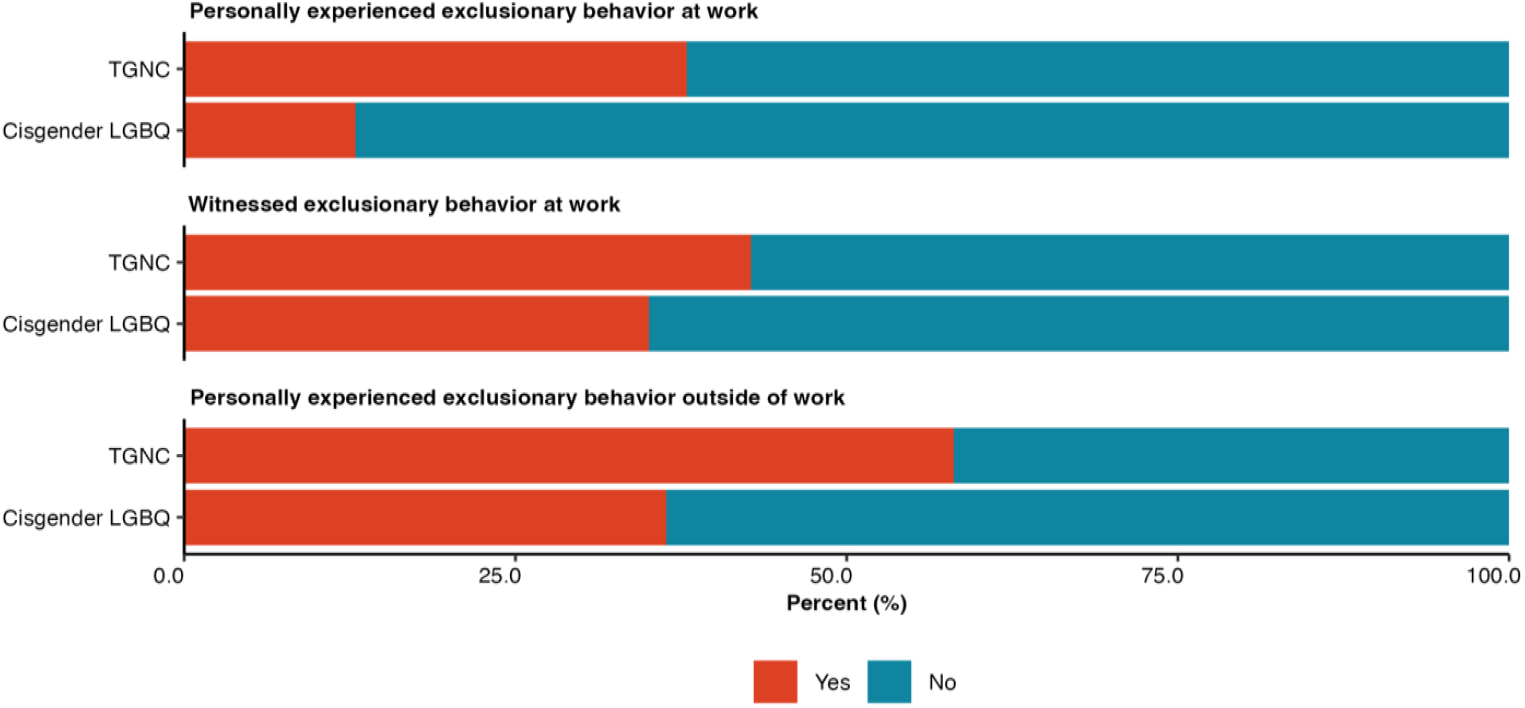
Distribution of responses from TGNC and cisgender LGBQ participants for whether they have (**A)** experienced exclusionary behavior at work, **(B)** witnessed exclusionary behavior at work, or **(C)** experienced exclusionary behavior outside of work.

### LGBTQ+ participants perceive moderate discrimination in biology

LGBTQ+ participants reported an average score of 3.2 out of 7 on the scale measuring discrimination against LGBTQ+ individuals in biology, where lower scores indicate greater discrimination in biology. When disaggregated by identity category, cisgender LGBQ participants scored an average of 3.5, while TGNC participants scored an average of 2.6. When controlling for race, professional position, and geographic location, TGNC participants perceived significantly more discrimination compared to cis LGBQ participants (β = -0.88, *p* < .0001). The distribution of responses across all LGBTQ+ participants is illustrated in Fig. 4 and full regression results in table S6.

**Figure 4.**
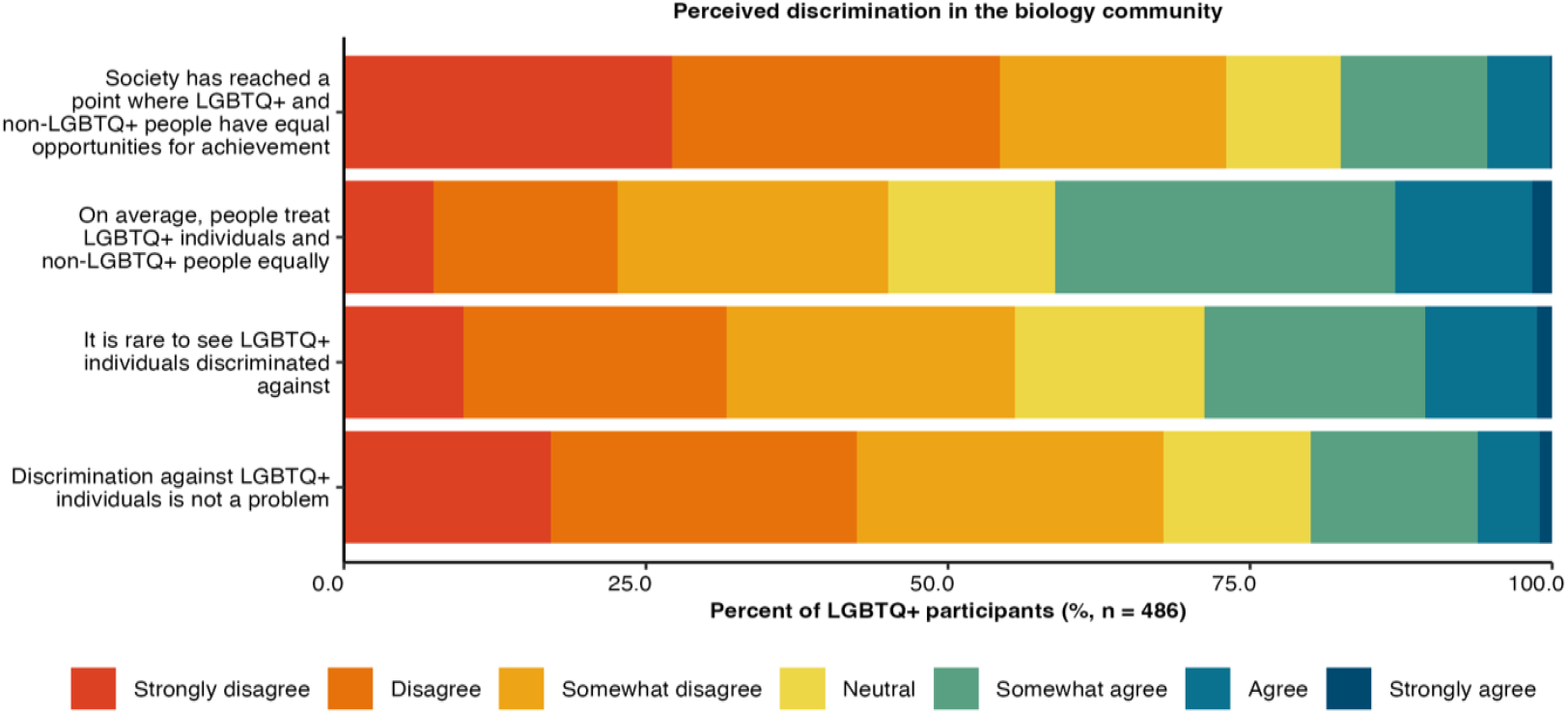
Distribution of responses from LGBTQ+ participants for the four indicated items assessing discrimination against LGBTQ+ individuals in biology.

### LGBTQ+ biologists are somewhat open about their identities to coworkers, but few are out to undergraduates

Over half of all LGBTQ+ participants were out to most people across an array of professional and personal contexts. Within professional contexts, participants were most likely to be out to postdocs or graduate students in their research groups (67.7%). Of the participants teaching graduate courses, 55.1% were out to graduate students in the course they teach and 16.2% were not. To coworkers or colleagues, 50.6% were out to all, while 5.4% were not to any. LGBTQ+ biologists were least out to undergraduates in their courses; 23.7% were not out to any undergraduates (Fig. 5A). Most LGBTQ+ biologists who were out to all coworkers or colleagues felt comfortable or very comfortable with their department climate (Fig. 5B).

**Figure 5.**
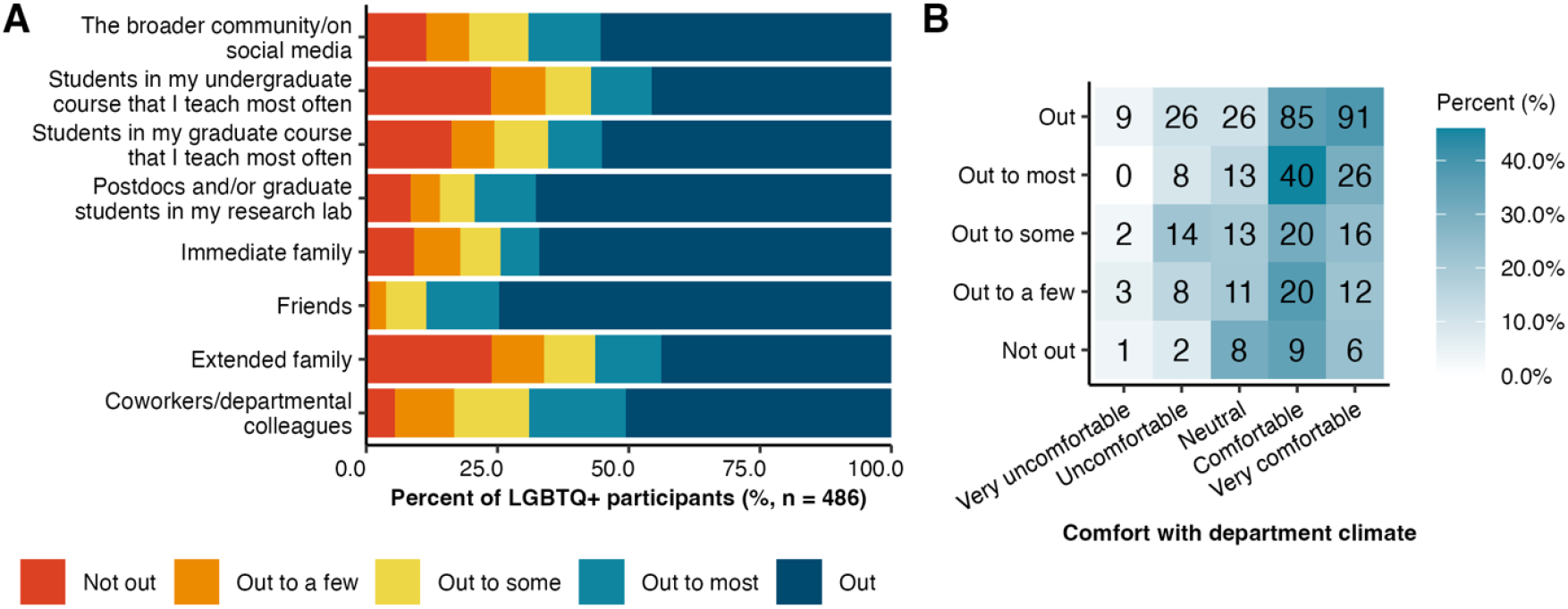
Distribution of responses from LGBTQ+ participants for **(A)** the degree to which they were out about their LGBTQ+ identities in various contexts, and **(B)** how outness to coworkers was related to comfort in the department/division. The numbers on the heatmap refer to the number of participants represented in that square (e.g., n=9 participants reported being very uncomfortable in their department and were out). The shade of the square represents the percent of LGBTQ+ participants represented per degree of outness (i.e., each row).

### LGBTQ+ biologists recommend increased institutional and social support

Participants were asked how their workplaces or professional societies could be more inclusive of LGBTQ+ employees and members. The three most common recommendations included: 1) communicating institutional support by speaking out for LGBTQ+ individuals, addressing LGBTQ-related issues directly, and using inclusive language (23.4%); 2) creating affinity groups or programming for LGBTQ+ individuals (16.4%); and 3) increasing LGBTQ+ visibility and representation (15.8%). Notably, many participants emphasized the need to raise awareness of issues facing TGNC biologists, such as ensuring access to gender-neutral bathrooms and using correct pronouns and names (13.4%). The full descriptions of these suggestions, including frequencies, are detailed in table S7.

## DISCUSSION

This study revealed remarkable differences between the experiences of cis LGBQ and TGNC individuals in biology, highlighting the need to disaggregate identities when examining the experiences of LGBTQ+ scientists. While cis LGBQ and non-LGBTQ+ biologists reported similar belonging and morale in the workplace and biology community, and similar comfort with the climate across a variety of professional contexts, TGNC biologists consistently reported lower belonging, morale, and climate than their straight and cis peers. Compared with data from 2021 showing that LGBTQ+ scientists are more likely to report career limitations and professional devaluation than non-LGBTQ+ scientists (2), the current data suggest that the increased public acceptance of orientation identities (29) combined with specific recommendations and efforts to increase inclusion among LGBTQ+ biologists (19, 30–32) may have resulted in an increase in belonging, comfort, and morale among LGBQ biologists; however, such efforts have fallen short of closing belonging, morale, and comfort gaps for their TGNC counterparts.

These gaps are unsurprising considering the overwhelming anti-TGNC legislation introduced within the U.S. in the past few years (33), and that the majority of research on queer biologists has been dominated by the experiences of cis LGBQ participants. However, recent studies are beginning to explicitly examine the experiences of TGNC people in biology, particularly undergraduates (27), and there is increased documentation of efforts to reform biology curriculum to be more inclusive of students with queer genders (26, 34). While these efforts are less likely to impact the experiences of practicing scientists, future evaluations are needed to assess whether they are effective in promoting inclusion of rising TGNC biologists.

The current study found that LGBTQ+ biologists are relatively open about their identities across contexts, and much more likely to be out than in prior years. Over half of LGBTQ+ biologists were out to all their colleagues, compared to only a third of 60 LGBTQ+ biologists in a 2019 study (22). While participants in the current study were least open about their identity in their undergraduate courses, with 24% not being out to any students, this is a vast improvement from the 2019 study where 65% of biologists were not out to any undergraduates. The rise in LGBTQ+ biologists coming out to undergraduates is encouraging, given recent research highlighting that most students in biology courses benefit from LGBTQ+ visibility and representation among scientists, regardless of whether they hold an LGBTQ+ identity (35, 36). While this progress is promising, the fact that nearly a quarter of LGBTQ+ biologists are not out to any undergraduates indicates ongoing challenges.

Given that over 20% of all LGBTQ+ participants and nearly 40% of TGNC participants experience intimidating, offensive, and/or hostile behavior at work due to their identity, there is an immediate need for workplaces and professional environments to evaluate their internal climates and work toward change as the biology community aims to become more diverse and inclusive. Recommendations from the LGBTQ+ participants are presented in Table S7 and include communicating institutional support for LGBTQ+ individuals and increasing programming to create more inclusive workplace environments. Employers interested in adopting these recommendations should explore resources, such as the Safe Zone Project (thesafezoneproject.com (37)), which provides curricula, activities and other resources for LGBTQ+ education sessions, and the Human Rights Campaign Establishing an Allies/Safe Zone Program (hrc.org/resources/establishing-an-allies-safe-zone-program (38)), which provides a step-by-step guide to establishing an LGBTQ+ ally program to make the culture of a workplace more inclusive of LGBTQ+ people.

The findings of this study point to critical areas of future research, particularly examining the intersectional experiences of individuals who hold specific identities within the LGBTQ+ umbrella and those who hold other marginalized identities. Great variation still exists among individuals within the subgroups of the LGBTQ+ community (39, 40)). The current study provides foundational insight into the current challenges experienced by cis LGBQ and TGNC biologists and invites the biology community to embrace the participant-generated recommendations in hopes that we can move toward a future where all biologists, regardless of their sexual orientation or gender identity, can thrive and contribute fully to the scientific enterprise.

Finally, the data for this study were collected in summer of 2023. Since January 2025, there has been a substantial increase in anti-LGBTQ+ and especially anti-trans rhetoric and policies in the US. This constantly changing political landscape may affect LGBTQ+ biologists’ perceptions of belonging and climate, as well as their experiences with harassment, emphasizing the importance of the continued assessment of LGBTQ+ biologists’ perceptions and experiences.

## MATERIALS AND METHODS

### Experimental design

To answer our research questions, we developed a survey with closed-ended and open-ended questions. A full copy of the survey questions is provided in Supplementary Materials.

### Sense of belonging and feelings of morale

We adapted the Course Cohesion Scale (41) to measure sense of belonging and feelings of morale associated with the biology community as well as participants’ professional societies and workplaces. The Course Cohesion Scale consists of two subscales – sense of belonging and feelings of moral – which each include 3 items. Participants respond on an 11-point Likert scale from strongly disagree (0) to strongly agree (10). We created aggregate scores for biology, professional society, and workplace sense of belonging and feelings of morale by calculating the mean of the three items. In our sample, we had excellent internal consistency for workplace sense of belonging (Cronbach’s α = 0.96), workplace feelings of morale (Cronbach’s α = 0.93), biology sense of belonging (Cronbach’s α = 0.96), biology feelings of morale (Cronbach’s α = 0.91), society sense of belonging (Cronbach’s α = 0.93-0.97), and society feelings of morale (Cronbach’s α = 0.90-0.93).

### Climate comfort

We assessed comfort with climate across professional and academic contexts using a single item per context. This item has previously been used in a LGBTQ+ climate survey from APS (17). Specifically, participants used a 5-point Likert scale from very uncomfortable to very comfortable to rate how comfortable they were with the climate in 1) their campus or company, 2) department or division, 3) research lab or group, 4) classroom where they are an instructor, and 5) classroom where they are a student.

### Demographic questions

Participants then answered a series of demographic questions with regard to their gender, race/ethnicity, LGBTQ+ identity, current professional position (e.g., graduate student, university faculty), and current region of residence.

### LGBTQ-specific questions

Participants who identified as a part of the LGBTQ+ community were asked a series of questions. First, we measured perceived bias against LGBTQ+ individuals in science by modifying a scale previously developed to assess bias against religious individuals in science (42). Participants rated four items on a scale from strongly disagree (1) to strongly agree (7). We created an aggregate score by taking the average of the four items. Our sample demonstrated good internal consistency (Cronbach’s α = 0.90).

Then, LGBTQ+ participants rated 20 items based on how well they described the atmosphere for LGBTQ+ employees at their workplace. These items have previously been used in a LGBTQ+ climate survey from APS (9, 17) and include 12 positively-worded inclusive climate items and 8 negatively-worded exclusion climate items. We created aggregate scores for both the inclusive climate and exclusive climate measures by taking the mean of the respective items. We found excellent internal consistency for these measures in our sample for both inclusion (Cronbach’s α = 0.93) and exclusion (Cronbach’s α = 0.91).

To determine the extent to which LGBTQ+ participants are open in various contexts, they used a 5-point scale from not out (1) to out (5) to describe the extent to which they typically revealed their LGBTQ+ identity to: friends, immediate family, extended family, coworkers, research group, graduate courses, undergraduate courses, and broader community. Participants had the option to select N/A if they did not interact with a particular group of individuals (e.g., they do not teach graduate courses).

Using a series of questions previously used in LGBTQ+ climate surveys in science (17), we assessed whether participants had experienced and/or witnessed LGBTQ+ exclusionary behavior. The survey concluded with two open-ended questions for participants to describe any ways in which their professional society and workplace could make them feel more included as an LGBTQ+ individual.

### Recruitment and distribution

To understand how the experiences of LGBTQ+ biologists compare to those of straight cis biologists, we aimed to recruit both LGBTQ+ and non-LGBTQ+ participants. Recruitment was initiated by the American Society of Cell Biology (ASCB) LGBTQ+ Committee. In summer 2023, ASCB leadership invited 5 societies to participate in helping to recruit biologists within their respective communities for this study. In addition to ASCB, 4 professional societies agreed to send out the study recruitment to their membership: Biophysical Society (BPS), Genetics Society of America (GSA), International Society for STEM Cell Research (ISSCR), and the Society for the Advancement of Biology Education Research (SABER). A snowball sampling approach was also used; the media teams from each community were encouraged to also use social media outlets to recruit any biologists interested in participating in the survey. Social media posts often encouraged biologists to invite other biologists to participate. The survey was open from June 1 to July 31 of 2023.

### Statistical analysis

We conducted all quantitative analyses in R. We report the percent of participants who selected a particular response for single-item measures and summarize responses by reporting the means for multi-item scales. For all outcomes, we report overall descriptive statistics for LGBTQ+ participants and then disaggregate the responses between TGNC participants and cisgender LGBQ participants in order to respect the differences in experiences for biologists across the LGBTQ+ spectrum. For all regression analyses, the reference group is cisgender straight participants. The specific analyses are organized by research question:

#### 1. To what extent does the belonging, morale, and comfort of LGBTQ+ and non-LGBTQ+ biologists differ?

To assess the extent to which LGBTQ+ individuals differ from non-LGBTQ+ individuals with regard to sense of belonging and feelings of morale across their professional societies, the biology community, and their workplace, we ran linear regressions using the stats package (*49*). We ran one set of regressions for orientation (cis LGBQ biologists compared to cisgender straight biologists) and another for gender (comparing TGNC biologists to cisgender straight biologists). For orientation, our predictors included whether the participant reported an LGBQ orientation identity (0/1), binary gender (cisgender man/cisgender woman), race (Asian, Latine, white, or other), professional position (university faculty, university staff, student, or outside of academia), and region (Midwest US, Northeast US, Southeast US, Southwest US, Western US, or outside of the US). We omitted participants who did not identify as cisgender in order to focus this analysis on the differences between cisgender biologists with or without LGBTQ+ orientation identities. Our reference groups were not reporting an LGBTQ+ orientation identity, man, white, outside of academia, and outside of the US. In our models examining the experiences of participants with LGBTQ+ gender identities, we modeled each of the outcomes by gender (TGNC/cisgender straight), race (Asian, Latine, white, other), professional position (university faculty, university staff, student, or outside of academia), and region (Midwest US, Northeast US, Southeast US, Southwest US, Western US, or outside of the US). Of note, cisgender LGBQ participants were not included in these analyses. Our reference groups were cisgender, white, outside of academia, and outside of the US. For the models assessing professional society sense of belonging and feelings of morale, we also included professional society as a predictor. In presenting the results, we have anonymized the society names and do not interpret any significant differences. If an individual was a member of multiple participating societies, they responded to these items for each society and each entry was included in the model. Thus, we included a random effect of participant to account for multiple responses from participants who were members of multiple societies. To account for multiple hypothesis testing across these six regressions, we used Bonferroni corrections and used 0.008 (0.05/6) as the *p*-value threshold.

Similarly, to assess the differences in experiences between LGBTQ+ and non-LGBTQ+ individuals with regard to their workplace climate, we ran two sets of ordinal regressions using the ordinal package. The two models had the same sets of predictors as described for the linear models above, with the outcome being participants’ responses to the question regarding how comfortable they are with the climate in various professional contexts (campus, department, lab, as a student, as an instructor). We again used Bonferroni corrections to account for multiple hypothesis testing and 0.01 (0.05/5) was the *p*-value threshold.

We calculated the variance inflation factor (VIF) for both sets of predictors using the car package, which indicated that there were no issues with multicollinearity among the predictors. Assumptions of linearity, homoscedasticity, and normality were checked and met for all linear regressions and proportional odds assumption was met for the ordinal regressions.

#### 2. How inclusive are biology workplaces for LGBTQ individuals?

We calculated the average score across the 12 positively worded items assessing LGBTQ+ inclusion and the average score across the 8 negatively worded LGBTQ+ items assessing exclusion. We also calculated these average scores for cisgender LGBQ participants as well as TGNC participants. We assessed potential statistically significant differences between TGNC participants and cisgender LGBQ participants using linear regressions for the inclusion and exclusion scales while controlling for race, professional position, and region.

#### 3. To what extent do LGBTQ+ biologists experience discrimination?

We assessed the percentage of TGNC individuals who personally experienced exclusionary behavior at work, personally witnessed exclusionary behavior at work, and personally experienced exclusionary behavior outside of work. This analysis was repeated for cis LGBQ individuals. We used binary logistic regression analyses to determine whether TGNC participants and cis LGBQ participants differed in their experiences with exclusionary behaviors while controlling for race, professional position, and region in the regressions pertaining to the workplace and race and region in the regression of experiencing exclusionary behavior outside of work. In reporting the results of the logistic regressions, we use the odds ratio (OR) to describe effect size. OR is calculated by exponentiating the β coefficient from the model output.

#### 4. To what extent are LGBTQ+ individuals out in different biology environments?

We calculated the percentages of all LGBTQ+ participants who were out in relation to eight circumstances. If a circumstance did not apply to an individual, they could select “N/A.” Percentages were calculated by dividing by the number of individuals who indicated the circumstance was applicable to them.

#### 5. What recommendations do LGBTQ+ biologists have for creating more inclusive biology environments?

To describe the recommendations participants had for making their professional societies and workplaces more inclusive for LGBTQ+ individuals, we used open-coding methods (43). Due to the parallel structure of the questions, we developed one codebook to analyze the responses. Two researchers (C.A.B. and P.B.B.) individually reviewed different randomly selected subsets of 10% of the responses and took analytic notes on themes that emerged from the text. Then, the researchers met to compare their codes and developed a preliminary codebook which included all of the themes that the researchers identified and ensured that all of the themes were distinct from each other. Then, one researcher (P.B.B.) tested the preliminary codebook on a randomly selected subset of 10% of the responses. Through this process, we ensured that no new codes were present in the responses and that our codebook was complete. The two researchers then independently coded the same subset of 15% of the responses using the codebook. Upon comparing their codes, they achieved excellent interrater reliability (Cohen’s K = 0.88) as evidence of the validity of the codebook. One researcher (P.B.B.) then coded all responses using the codebook. We report the percent of participants who included each theme in their response as well as example quotes for the two prompts in table S7.

## ACKNOLWEDGEMENTS

We would like to acknowledge the leaders and communication staff of the professional societies who generously supported this work by helping to recruit individuals within their respective communities: Jennifer Pesanelli & Laura Phelan (BPS), Tracey DePellegrin, Sarah Bay, & Sana Hussain (GSA), Keith Alm & Kym Kilbourne (ISSCR), and Miriam Segura, Kelsey Metzger, Melinda Owens, Sheela Vemu & Emily Grunspan (SABER).

