## Supplemental Material for "LGBTQ+ realities in the biological sciences"

**This PDF file includes:**

Supporting text
Tables S1 to S7
Legends for Datasets S1 to S2

**Supporting Information Text**

Copy of survey questions

Please select which of the following professional societies you are associated with. You do not have to be an active member to be associated with a particular society.

- American Society of Cell Biology (ASCB)
- Biophysical Society (BPS)
- Genetics Society of America (GSA)
- International Society for Stem Cell Research (ISSCR)
- Society for the Advancement of Biology Education Research (SABER)
- I am not currently associated with any of the above societies

*Based on society(ies) selected in previous question:*
Please answer the following questions about **the [society] community.** *Answered on a scale from 0 (strongly disagree) to 10 (strongly agree)*

1. I feel a sense of belonging to [society]
2. I feel that I am a member of the [society] community
3. I see myself as part of the [society] community
4. I am enthusiastic about [society]
5. I am happy to be associated with [society]
6. [Society] is one of the best professional societies

Please answer the following questions about **your department or workplace.** *Answered on a scale from 0 (strongly disagree) to 10 (strongly agree)*

1. I feel a sense of belonging to my workplace
2. I feel that I am a member of my workplace community
3. I see myself as part of my workplace community
4. I am enthusiastic about where I work
5. I am happy to be at my place of work
6. I am at one of the best places to work

What subdiscipline(s) of biology do you work within?

- Anatomy
- Biochemistry
- Bioinformatics
- Biophysics
- Biotechnology
- Botany
- Cell biology
- Developmental biology
- Discipline-based education (DBER)
- Ecology
- Embryology
- Entomology
- Genetics
- Immunology
- Marine biology
- Microbiology
- Molecular biology
- Molecular genetics
- Mycology
- Neuroscience
- Parasitology
- Pathology
- Physiology
- Systems biology
- Zoology
- Other, please describe
- I do not identify as working within a sub-discipline of biology *sent to end of survey*

Please answer the following questions about **the broader biology community.** The broader biology community includes everyone you know in biology. *Answered on a scale from 0 (strongly disagree) to 10 (strongly agree)*

1. I feel a sense of belonging to the biology community
2. I feel that I am a member of the biology community
3. I see myself as part of the biology community
4. I am enthusiastic about the biology community
5. I am happy to be associated with the biology community
6. The biology community is one of the best scientific communities

Climate is defined as the "current attitudes, behaviors, and standards held by faculty, staff, and students concerning access for, inclusion of, and level of respect for individual and group needs, abilities and potential" (APS, 2015).

Overall how comfortable are you with the climate in the following areas? *Answered on a scale from very uncomfortable (1) to very comfortable (5), with the option to select N/A does not apply to me*

1. Campus or company
2. Department or division
3. Research lab or group
4. Classroom where I am an instructor
5. Classroom where I am a student

Next, we ask a series of demographic questions to better understand the identities of those in the biology community. You are always welcome to skip questions or decline to state if you would rather not answer.

I most closely identify as

- Woman
- Man
- Nonbinary or genderfluid
- Agender
- A gender that is not listed ______________________________
- Decline to state

I most closely identify as

- American Indian or Alaska Native
- Asian
- Black or African American
- Hispanic, Latin*, or Spanish origin
- Pacific Islander
- White
- More than one race/ethnicity
- Other (please describe) ______________________________
- Decline to state

Do you identify as a member of the LGBTQ+ community?

- Yes
- No
- Decline to state

*If yes:* Please select the word or words that best describe your LGBTQ+ identities referring to sexuality and/or romantic attraction.

- Lesbian or Gay
- Bisexual
- Queer as it relates to my sexuality
- Asexual/aromantic
- Pansexual
- Questioning
- An identity not listed, please describe
  ________________________________________________
- Straight or I do not identify within the LGBTQ+ community as it relates to my sexuality or romantic attraction
- Decline to state

Please select the word or words that best describe your LGBTQ+ identities referring to gender identity.

- Transgender man
- Transgender woman
- Gender-queer or gender non-binary
- Agender
- Questioning
- A gender that is not listed, please describe
  ________________________________________________
- Cis-gender or I do not identify within the LGBTQ+ community as it relates to my gender identity
- Decline to state

I most closely identify as

- An undergraduate student
- A graduate student
- A postdoctoral scholar
- An assistant professor
- An associate professor
- A full professor
- A lecture or instructor
- A non-tenure track faculty member or research staff
- Employed in a government position
- Employed in an industry position
- Other, please describe ________________________________________________
- Decline to state

What type of academic institution do you identify with?

- I do not work at an academic institution
- R1 doctoral university
- R2 doctoral university
- R3 doctoral university
- Master's college or university
- Baccalaureate college or university
- Associates college or community college
- Tribal college or university
- Decline to state

In which country do you currently reside? *Dropdown selection*

*If in the USA:* Where do you currently reside?

- Western US
- Southwestern US
- Southeastern US
- Northeastern US
- Midwestern US
- Decline to state

*If identify as LGBTQ+:* The following questions are intended to assess how inclusive the biology community is of LGBTQ+ individuals. Please answer the following questions about your experience. If you prefer not to answer, you may skip any questions.

*Answered on a scale from strongly disagree (1) to strongly agree (7)*

1. Discrimination against LGBTQ+ individuals is not a problem in the biology community
2. It is rare to see LGBTQ+ individuals discriminated against in the biology community
3. On average, people in the biology community treat LGBTQ+ individuals and non-LGBTQ+ people equally
4. Society has reached a point where LGBTQ+ and non-LGBTQ+ people have equal opportunities for achievement in the biology community

Please rate the following items according to how well they describe the atmosphere for employees who identify within the LGBTQ+ community in your department/workplace. *Answered on a scale from strongly disagree (1) to strongly agree (5)*

1. LGBTQ+ employees are treated with respect
2. LGBTQ+ employees must be secretive
3. Coworkers are as likely to ask nice, interested questions about LGBTQ+ relationships as they are about non-LGBTQ+ relationships
4. LGBTQ+ people consider it a comfortable place to work
5. Non-LGBTQ+ employees are comfortable engaging in LGBTQ+ friendly humor with LGBTQ+ employees (for example, kidding them about a date)
6. The atmosphere for LGBTQ+ employees is oppressive
7. LGBTQ+ employees feel accepted by coworkers
8. Coworkers make comments that seem to indicate a lack of awareness of LGBTQ+ issues
9. Employees are expected to not act "too gay"
10. LGBTQ+ employees fear job loss because of their identity
11. My immediate work group is supportive of LGBTQ+ coworkers
12. LGBTQ+ employees are comfortable talking about their personal lives with coworkers
13. There is pressure for LGBTQ+ employees to stay closeted (to conceal their sexual orientation or gender identity/expression)
14. Employee LGBTQ+ identity does not seem to be an issue
15. LGBTQ+ employees are met with thinly veiled hostility (for example, scornful looks or icy tone of voice)
16. The company or institution as a whole provides a supportive environment for LGBTQ+ people
17. LGBTQ+ employees are free to be themselves
18. LGBTQ+ people are less likely to be mentored
19. LGBTQ+ employees feel free to display pictures of their partners
20. The atmosphere for LGBTQ+ employees is improving

To what extent do you typically reveal your LGBTQ+ identity in the following contexts. *Answered on a scale from not out (1) to out (5) with the option to select N/A I do not interact with these individuals*

1. Friends
2. Immediate family
3. Extended family
4. Coworkers/departmental colleagues
5. To postdocs and/or graduate students in my research lab
6. To students in my graduate course that I teach most often
7. To students in my undergraduate course that I teach most often
8. To the broader community/on social media

Within the past year, have **you personally** experienced any exclusionary (e.g., shunned, ignored), intimidating, offensive and/or hostile conduct (harassing behavior) because of your LGBTQ+ identity that has interfered with your ability to work or learn on your department or workplace?

- Yes
- No
- Decline to state

Within the past year, have you observed or personally been made aware of any conduct directed **toward a person or group of people on campus or at your department or workplace** that you believe has created an exclusionary (e.g. shunned, ignored), intimidating, offensive, and/or hostile (harassing) working or learning environment because of their LGBTQ+ identity?

- Yes
- No
- Decline to state

Within the past year, have **you personally** experienced any exclusionary (e.g., shunned, ignored), intimidating, offensive and/or hostile conduct (harassing behavior) **outside of your department or workplace** because of your LGBTQ+ identity?

- Yes
- No
- Decline to state

What, if anything, could your respective **professional society/societies** do to make you feel more included as an LGBTQ+ individual? *Open-ended*

What, if anything, could your **workplace** do to make you feel more included as an LGBTQ+ individual? *Open-ended*

**Table S1.** Participant demographic characteristics.

| | Demographic | Percent (n) |
|---|---|---|
| Gender | Woman | 47.0 (667) |
| | Man | 41.3 (586) |
| | Non-Binary Or Gender Fluid | 5.1 (72) |
| | Agender | 0.9 (13) |
| | A Gender That Is Not Listed | 0.8 (11) |
| | Decline To State | 4.9 (70) |
| Race/ ethnicity | White | 64.5 (915) |
| | Asian | 11.5 (163) |
| | Hispanic, Latin*, Or Spanish Origin | 8.2 (116) |
| | More Than One Race/Ethnicity | 3.7 (53) |
| | Identity(ies) Not Listed | 3.4 (48) |
| | Black Or African American | 2.6 (37) |
| | American Indian Or Alaska Native | 0.6 (8) |
| | Pacific Islander | 0.1 (2) |
| | Decline To State | 5.4 (77) |
| LGBTQ+ | No | 58.3 (827) |
| | Yes | 34.2 (486) |
| | Decline To State | 7.5 (106) |
| LGBTQ+ identity group | Cisgender LGBQ | 22.1 (313) |
| | Trans, nonbinary, genderfluid, or genderqueer (TGNC) | 10.8 (153) |
| | Decline To State | 1.4 (20) |
| LGBTQ+ gender identity (can select >1) | Agender | 1.3 (19) |
| | Gender-queer or gender non-binary | 7.8 (111) |
| | Questioning | 1.8 (26) |
| | Transgender man | 1.3 (18) |
| | Transgender woman | 0.7 (10) |
| | A gender not listed | 1.3 (18) |
| | Decline to state | 0.8 (11) |

| | | |
|---|---|---|
| LGBTQ+ orientation identity (can select >1) | Asexual or aromantic | 2.7 (38) |
| | Bisexual | 9.7 137) |
| | Lesbian or gay | 18.9 (268) |
| | Pansexual | 3.5 (49) |
| | Queer | 10.5 (149) |
| | Questioning | 0.4 (6) |
| | An identity not listed | 0.6 (9) |
| | Decline to state | 0.2 (3) |
| Geographic location | Outside of US | 22.1 (314) |
| | Northeast US | 19.7 (280) |
| | Midwest US | 15.4 (219) |
| | West US | 14.7 (209) |
| | Southeast US | 12.8 (181) |
| | Southwest US | 5.6 (80) |
| | Decline to state | 9.6 (136) |
| Professional position | University faculty | 49.9 (708) |
| | University staff, lecturer, or postdoc | 20.4 (290) |
| | University undergrad or grad student | 15.3 (217) |
| | Position outside of academia (e.g., govt. or industry, nonprofit) | 11.0 (156) |
| | Decline to state | 3.4 (48) |
| Society (can select >1) | ASCB | 416 |
| | BPS | 172 |
| | GSA | 353 |
| | ISSCR | 162 |
| | SABER | 176 |
| | None | 375 |

**Table S2.** Results from linear regressions for sense of belonging and feelings of morale in the workplace, biology community, and professional societies.

| **Outcome** | **Predictor** | **beta** | **SE** | **p-value** |
|---|---|---|---|---|
| Workplace sense of belonging | (Intercept) | 8.018 | 0.267 | **0.000** |
| | Cis LGBQ (yes) | -0.351 | 0.166 | 0.034 |
| | Gender (cis woman) | -0.194 | 0.144 | 0.178 |
| | Race (Asian) | -0.081 | 0.232 | 0.726 |
| | Race (Latinx) | -0.277 | 0.261 | 0.290 |
| | Race (other) | -0.015 | 0.239 | 0.950 |
| | Position (faculty) | -0.064 | 0.235 | 0.784 |
| | Position (university staff) | -0.500 | 0.262 | 0.057 |
| | Position (student) | -0.334 | 0.290 | 0.250 |
| | Region (Midwest US) | -0.422 | 0.225 | 0.061 |
| | Region (NE US) | 0.049 | 0.214 | 0.819 |
| | Region (SE US) | 0.142 | 0.237 | 0.549 |
| | Region (SW US) | -0.251 | 0.321 | 0.435 |
| | Region (West US) | 0.281 | 0.230 | 0.222 |
| Workplace feelings of morale | (Intercept) | 7.685 | 0.279 | **0.000** |
| | Cis LGBQ (yes) | -0.181 | 0.173 | 0.295 |
| | Gender (cis woman) | -0.313 | 0.150 | 0.037 |
| | Race (Asian) | 0.011 | 0.242 | 0.965 |
| | Race (Latinx) | -0.048 | 0.273 | 0.859 |
| | Race (other) | 0.062 | 0.250 | 0.805 |
| | Position (faculty) | -0.436 | 0.245 | 0.075 |
| | Position (university staff) | -0.289 | 0.274 | 0.291 |
| | Position (student) | -0.521 | 0.303 | 0.086 |
| | Region (Midwest US) | -0.770 | 0.235 | **0.001** |
| | Region (NE US) | -0.164 | 0.223 | 0.464 |
| | Region (SE US) | -0.042 | 0.247 | 0.866 |
| | Region (SW US) | -0.277 | 0.336 | 0.409 |
| | Region (West US) | 0.527 | 0.240 | 0.028 |
| Biology sense of belonging | (Intercept) | 7.722 | 0.236 | **0.000** |
| | Cis LGBQ (yes) | -0.157 | 0.146 | 0.282 |
| | Gender (cis woman) | -0.255 | 0.126 | 0.044 |
| | Race (Asian) | 0.099 | 0.204 | 0.627 |
| | Race (Latinx) | 0.231 | 0.229 | 0.313 |
| | Race (other) | -0.055 | 0.210 | 0.795 |
| | Position (faculty) | 0.122 | 0.207 | 0.554 |
| | Position (university staff) | -0.091 | 0.231 | 0.694 |
| | Position (student) | -0.332 | 0.255 | 0.194 |
| | Region (Midwest US) | -0.215 | 0.198 | 0.278 |
| | Region (NE US) | -0.121 | 0.188 | 0.518 |
| | Region (SE US) | -0.040 | 0.208 | 0.846 |
| | Region (SW US) | -0.339 | 0.282 | 0.230 |
| | Region (West US) | 0.083 | 0.202 | 0.681 |
| Biology feelings of morale | (Intercept) | 7.729 | 0.212 | **0.000** |
| | Cis LGBQ (yes) | -0.161 | 0.131 | 0.220 |

| | | | | |
|---|---|---|---|---|
| | Gender (cis woman) | -0.133 | 0.113 | 0.241 |
| | Race (Asian) | 0.033 | 0.182 | 0.857 |
| | Race (Latinx) | 0.018 | 0.205 | 0.932 |
| | Race (other) | 0.160 | 0.188 | 0.396 |
| | Position (faculty) | 0.107 | 0.185 | 0.565 |
| | Position (university staff) | -0.034 | 0.207 | 0.868 |
| | Position (student) | 0.208 | 0.229 | 0.362 |
| | Region (Midwest US) | -0.228 | 0.177 | 0.199 |
| | Region (NE US) | 0.075 | 0.168 | 0.657 |
| | Region (SE US) | 0.229 | 0.186 | 0.219 |
| | Region (SW US) | -0.190 | 0.253 | 0.452 |
| | Region (West US) | 0.342 | 0.181 | 0.059 |
| Workplace sense of belonging | (Intercept) | 8.026 | 0.290 | **0.000** |
| | TGNC (yes) | -1.002 | 0.232 | **0.000** |
| | Race (Asian) | -0.095 | 0.241 | 0.694 |
| | Race (Latinx) | -0.238 | 0.305 | 0.436 |
| | Race (other) | -0.014 | 0.248 | 0.954 |
| | Position (faculty) | -0.366 | 0.264 | 0.166 |
| | Position (university staff) | -0.795 | 0.300 | 0.008 |
| | Position (student) | -0.306 | 0.334 | 0.359 |
| | Region (Midwest US) | -0.261 | 0.249 | 0.295 |
| | Region (NE US) | 0.317 | 0.234 | 0.176 |
| | Region (SE US) | 0.303 | 0.277 | 0.274 |
| | Region (SW US) | -0.221 | 0.372 | 0.553 |
| | Region (West US) | 0.385 | 0.255 | 0.131 |
| Workplace feelings of morale | (Intercept) | 7.534 | 0.306 | **0.000** |
| | TGNC (yes) | -0.706 | 0.244 | **0.004** |
| | Race (Asian) | -0.035 | 0.254 | 0.891 |
| | Race (Latinx) | -0.109 | 0.322 | 0.736 |
| | Race (other) | 0.146 | 0.262 | 0.578 |
| | Position (faculty) | -0.577 | 0.278 | 0.038 |
| | Position (university staff) | -0.390 | 0.316 | 0.217 |
| | Position (student) | -0.277 | 0.352 | 0.431 |
| | Region (Midwest US) | -0.597 | 0.262 | 0.023 |
| | Region (NE US) | 0.118 | 0.246 | 0.632 |
| | Region (SE US) | -0.010 | 0.292 | 0.974 |
| | Region (SW US) | -0.348 | 0.392 | 0.375 |
| | Region (West US) | 0.359 | 0.269 | 0.183 |
| Biology sense of belonging | (Intercept) | 7.522 | 0.254 | **0.000** |
| | TGNC (yes) | -1.052 | 0.201 | **0.000** |
| | Race (Asian) | 0.253 | 0.209 | 0.227 |
| | Race (Latinx) | 0.235 | 0.264 | 0.373 |
| | Race (other) | -0.115 | 0.216 | 0.593 |
| | Position (faculty) | -0.105 | 0.231 | 0.648 |
| | Position (university staff) | -0.234 | 0.261 | 0.370 |
| | Position (student) | -0.325 | 0.291 | 0.264 |
| | Region (Midwest US) | -0.016 | 0.216 | 0.940 |
| | Region (NE US) | 0.397 | 0.203 | 0.051 |
| | Region (SE US) | 0.325 | 0.241 | 0.177 |
| | Region (SW US) | -0.493 | 0.321 | 0.125 |

| | | | | |
|---|---|---|---|---|
| | Region (West US) | 0.197 | 0.222 | 0.375 |
| Biology feelings of morale | (Intercept) | 7.644 | 0.233 | **0.000** |
| | TGNC (yes) | -1.021 | 0.185 | **0.000** |
| | Race (Asian) | 0.174 | 0.192 | 0.366 |
| | Race (Latinx) | -0.012 | 0.242 | 0.960 |
| | Race (other) | -0.016 | 0.198 | 0.938 |
| | Position (faculty) | -0.091 | 0.212 | 0.668 |
| | Position (university staff) | -0.282 | 0.240 | 0.241 |
| | Position (student) | 0.145 | 0.267 | 0.587 |
| | Region (Midwest US) | -0.073 | 0.199 | 0.713 |
| | Region (NE US) | 0.515 | 0.187 | **0.006** |
| | Region (SE US) | 0.592 | 0.221 | **0.008** |
| | Region (SW US) | -0.233 | 0.295 | 0.430 |
| | Region (West US) | 0.417 | 0.204 | 0.041 |
| Society sense of belonging | (Intercept) | 6.963 | 0.311 | **0.000** |
| | Cis LGBQ (yes) | -0.398 | 0.203 | 0.050 |
| | Gender (cis woman) | -0.354 | 0.138 | 0.011 |
| | Race (Asian) | 0.158 | 0.213 | 0.457 |
| | Race (Latinx) | -0.011 | 0.259 | 0.966 |
| | Race (other) | -0.168 | 0.219 | 0.443 |
| | Society B | 0.860 | 0.220 | **0.000** |
| | Society C | 0.371 | 0.176 | 0.035 |
| | Society D | -0.276 | 0.240 | 0.251 |
| | Society E | 0.103 | 0.227 | 0.652 |
| | Position (faculty) | 0.055 | 0.245 | 0.822 |
| | Position (university staff) | -0.166 | 0.276 | 0.549 |
| | Position (student) | 0.065 | 0.334 | 0.845 |
| | Region (Midwest US) | 0.259 | 0.230 | 0.261 |
| | Region (NE US) | 0.272 | 0.213 | 0.202 |
| | Region (SE US) | 0.517 | 0.246 | 0.036 |
| | Region (SW US) | 0.266 | 0.322 | 0.408 |
| | Region (West US) | 0.340 | 0.234 | 0.146 |
| Society feelings of morale | (Intercept) | 7.859 | 0.266 | **0.000** |
| | Cis LGBQ (yes) | -0.500 | 0.174 | **0.004** |
| | Gender (cis woman) | -0.080 | 0.118 | 0.501 |
| | Race (Asian) | 0.032 | 0.182 | 0.862 |
| | Race (Latinx) | -0.088 | 0.222 | 0.692 |
| | Race (other) | -0.021 | 0.187 | 0.912 |
| | Society B | 0.317 | 0.188 | 0.092 |
| | Society C | -0.006 | 0.151 | 0.968 |
| | Society D | -0.176 | 0.206 | 0.392 |
| | Society E | -0.080 | 0.195 | 0.680 |
| | Position (faculty) | 0.044 | 0.210 | 0.835 |
| | Position (university staff) | 0.026 | 0.237 | 0.913 |
| | Position (student) | 0.019 | 0.286 | 0.947 |
| | Region (Midwest US) | -0.053 | 0.197 | 0.787 |
| | Region (NE US) | 0.134 | 0.183 | 0.463 |
| | Region (SE US) | 0.316 | 0.211 | 0.135 |
| | Region (SW US) | 0.213 | 0.276 | 0.440 |
| | Region (West US) | 0.115 | 0.200 | 0.566 |

| | | | | |
|---|---|---|---|---|
| Society sense of belonging | (Intercept) | 6.628 | 0.313 | **0.000** |
| | TGNC (yes) | -1.077 | 0.289 | **0.000** |
| | Race (Asian) | 0.168 | 0.210 | 0.424 |
| | Race (Latinx) | 0.057 | 0.288 | 0.842 |
| | Race (other) | -0.249 | 0.223 | 0.266 |
| | Society B | 0.918 | 0.223 | **0.000** |
| | Society C | 0.379 | 0.180 | 0.036 |
| | Society D | -0.179 | 0.248 | 0.472 |
| | Society E | 0.204 | 0.239 | 0.393 |
| | Position (faculty) | 0.172 | 0.256 | 0.502 |
| | Position (university staff) | -0.130 | 0.291 | 0.654 |
| | Position (student) | 0.116 | 0.363 | 0.749 |
| | Region (Midwest US) | 0.271 | 0.230 | 0.241 |
| | Region (NE US) | 0.447 | 0.215 | 0.038 |
| | Region (SE US) | 0.518 | 0.258 | 0.045 |
| | Region (SW US) | 0.263 | 0.342 | 0.442 |
| | Region (West US) | 0.200 | 0.235 | 0.396 |
| Society feelings of morale | (Intercept) | 7.674 | 0.270 | **0.000** |
| | TGNC (yes) | -0.931 | 0.250 | **0.000** |
| | Race (Asian) | 0.053 | 0.181 | 0.770 |
| | Race (Latinx) | 0.036 | 0.249 | 0.886 |
| | Race (other) | -0.021 | 0.193 | 0.915 |
| | Society B | 0.413 | 0.192 | 0.032 |
| | Society C | 0.025 | 0.155 | 0.873 |
| | Society D | -0.064 | 0.214 | 0.764 |
| | Society E | -0.003 | 0.206 | 0.989 |
| | Position (faculty) | 0.112 | 0.220 | 0.613 |
| | Position (university staff) | -0.018 | 0.251 | 0.942 |
| | Position (student) | 0.041 | 0.313 | 0.896 |
| | Region (Midwest US) | -0.064 | 0.199 | 0.749 |
| | Region (NE US) | 0.332 | 0.186 | 0.074 |
| | Region (SE US) | 0.297 | 0.223 | 0.182 |
| | Region (SW US) | 0.201 | 0.295 | 0.496 |
| | Region (West US) | 0.126 | 0.203 | 0.533 |

Reference groups are straight, cisgender, white, outside of academia, outside of the US, and Society A. Bold text represents statistical significance after Bonferroni corrections.

**Table S3.** Results from ordinal regressions for climate across professional and academic contexts.

| Context | Predictor | beta | SE | OR | p-value |
|---|---|---|---|---|---|
| Campus | Cis LGBQ (yes) | -0.205 | 0.137 | 0.815 | 0.135 |
| | Gender (cis woman) | -0.320 | 0.120 | 0.726 | **0.008** |
| | Race (Asian) | -0.153 | 0.192 | 0.858 | 0.427 |
| | Race (Latinx) | -0.066 | 0.213 | 0.937 | 0.758 |
| | Race (other) | -0.358 | 0.199 | 0.699 | 0.072 |
| | Position (faculty) | -0.480 | 0.240 | 0.619 | 0.046 |
| | Position (university staff) | -0.493 | 0.258 | 0.611 | 0.056 |
| | Position (student) | -0.367 | 0.274 | 0.693 | 0.181 |
| | Region (Midwest US) | -0.001 | 0.185 | 0.999 | 0.996 |
| | Region (NE US) | 0.392 | 0.179 | 1.479 | 0.028 |
| | Region (SE US) | 0.123 | 0.194 | 1.131 | 0.526 |
| | Region (SW US) | -0.225 | 0.263 | 0.798 | 0.392 |
| | Region (West US) | 0.583 | 0.195 | 1.791 | **0.003** |
| Department | Cis LGBQ (yes) | -0.020 | 0.137 | 0.980 | 0.885 |
| | Gender (cis woman) | -0.247 | 0.121 | 0.781 | 0.041 |
| | Race (Asian) | -0.181 | 0.194 | 0.835 | 0.351 |
| | Race (Latinx) | -0.167 | 0.214 | 0.846 | 0.435 |
| | Race (other) | -0.450 | 0.199 | 0.638 | 0.024 |
| | Position (faculty) | -0.389 | 0.244 | 0.677 | 0.110 |
| | Position (university staff) | -0.761 | 0.259 | 0.467 | **0.003** |
| | Position (student) | -0.691 | 0.276 | 0.501 | 0.012 |
| | Region (Midwest US) | 0.199 | 0.186 | 1.220 | 0.285 |
| | Region (NE US) | 0.419 | 0.178 | 1.521 | 0.018 |
| | Region (SE US) | 0.487 | 0.197 | 1.627 | 0.014 |
| | Region (SW US) | -0.284 | 0.261 | 0.753 | 0.277 |
| | Region (West US) | 0.408 | 0.194 | 1.503 | 0.036 |
| Lab | Cis LGBQ (yes) | -0.003 | 0.170 | 0.997 | 0.985 |
| | Gender (cis woman) | -0.174 | 0.150 | 0.840 | 0.245 |
| | Race (Asian) | -0.602 | 0.218 | 0.548 | **0.006** |
| | Race (Latinx) | -0.409 | 0.252 | 0.664 | 0.104 |
| | Race (other) | -0.333 | 0.247 | 0.717 | 0.178 |
| | Position (faculty) | 0.628 | 0.283 | 1.874 | 0.027 |
| | Position (university staff) | -0.330 | 0.295 | 0.719 | 0.264 |
| | Position (student) | -0.332 | 0.313 | 0.718 | 0.289 |
| | Region (Midwest US) | 0.353 | 0.225 | 1.424 | 0.117 |
| | Region (NE US) | 0.674 | 0.219 | 1.962 | **0.002** |
| | Region (SE US) | 0.748 | 0.245 | 2.113 | **0.002** |
| | Region (SW US) | 0.451 | 0.324 | 1.570 | 0.165 |
| | Region (West US) | 0.650 | 0.240 | 1.915 | **0.007** |
| Teaching | Cis LGBQ (yes) | -0.266 | 0.175 | 0.766 | 0.128 |
| | Gender (cis woman) | -0.397 | 0.151 | 0.673 | **0.009** |
| | Race (Asian) | -0.802 | 0.243 | 0.449 | **0.001** |
| | Race (Latinx) | -0.157 | 0.281 | 0.855 | 0.577 |
| | Race (other) | -0.459 | 0.254 | 0.632 | 0.070 |
| | Position (faculty) | 1.194 | 0.556 | 3.301 | 0.032 |

| | | | | | |
|---|---|---|---|---|---|
| | Position (university staff) | 0.622 | 0.573 | 1.862 | 0.278 |
| | Position (student) | 0.689 | 0.585 | 1.991 | 0.239 |
| | Region (Midwest US) | 0.621 | 0.240 | 1.861 | **0.010** |
| | Region (NE US) | 0.703 | 0.226 | 2.021 | **0.002** |
| | Region (SE US) | 0.468 | 0.237 | 1.597 | 0.048 |
| | Region (SW US) | 0.547 | 0.339 | 1.728 | 0.107 |
| | Region (West US) | 0.804 | 0.255 | 2.235 | **0.002** |
| Student | Cis LGBQ (yes) | -0.542 | 0.319 | 0.581 | 0.089 |
| | Gender (cis woman) | -0.581 | 0.288 | 0.559 | 0.044 |
| | Race (Asian) | -0.714 | 0.411 | 0.490 | 0.082 |
| | Race (Latinx) | 0.772 | 0.437 | 2.164 | 0.077 |
| | Race (other) | -0.851 | 0.451 | 0.427 | 0.059 |
| | Position (faculty) | -0.081 | 0.696 | 0.922 | 0.907 |
| | Position (university staff) | -0.844 | 0.694 | 0.430 | 0.224 |
| | Position (student) | -0.345 | 0.665 | 0.708 | 0.604 |
| | Region (Midwest US) | 0.987 | 0.437 | 2.684 | 0.024 |
| | Region (NE US) | 1.447 | 0.541 | 4.251 | **0.007** |
| | Region (SE US) | 1.241 | 0.422 | 3.457 | **0.003** |
| | Region (SW US) | 1.526 | 0.744 | 4.602 | 0.040 |
| | Region (West US) | 0.798 | 0.425 | 2.221 | 0.060 |
| Campus | TGNC (yes) | -0.839 | 0.188 | 0.432 | **0.000** |
| | Race (Asian) | -0.128 | 0.194 | 0.880 | 0.510 |
| | Race (Latinx) | 0.078 | 0.244 | 1.081 | 0.748 |
| | Race (other) | -0.127 | 0.202 | 0.881 | 0.531 |
| | Position (faculty) | -0.552 | 0.270 | 0.576 | 0.041 |
| | Position (university staff) | -0.523 | 0.290 | 0.593 | 0.071 |
| | Position (student) | -0.261 | 0.313 | 0.770 | 0.404 |
| | Region (Midwest US) | 0.138 | 0.199 | 1.148 | 0.488 |
| | Region (NE US) | 0.570 | 0.189 | 1.769 | **0.003** |
| | Region (SE US) | 0.391 | 0.226 | 1.478 | 0.084 |
| | Region (SW US) | -0.249 | 0.303 | 0.780 | 0.412 |
| | Region (West US) | 0.514 | 0.208 | 1.672 | 0.014 |
| Department | TGNC (yes) | -0.576 | 0.186 | 0.562 | **0.002** |
| | Race (Asian) | -0.174 | 0.196 | 0.841 | 0.375 |
| | Race (Latinx) | -0.145 | 0.241 | 0.865 | 0.549 |
| | Race (other) | -0.349 | 0.203 | 0.705 | 0.086 |
| | Position (faculty) | -0.451 | 0.281 | 0.637 | 0.109 |
| | Position (university staff) | -0.805 | 0.299 | 0.447 | **0.007** |
| | Position (student) | -0.631 | 0.321 | 0.532 | 0.049 |
| | Region (Midwest US) | 0.328 | 0.202 | 1.388 | 0.105 |
| | Region (NE US) | 0.506 | 0.188 | 1.659 | **0.007** |
| | Region (SE US) | 0.560 | 0.227 | 1.751 | 0.013 |
| | Region (SW US) | -0.191 | 0.297 | 0.827 | 0.521 |
| | Region (West US) | 0.294 | 0.210 | 1.342 | 0.161 |
| Lab | TGNC (yes) | -0.445 | 0.216 | 0.641 | 0.040 |
| | Race (Asian) | -0.719 | 0.219 | 0.487 | **0.001** |
| | Race (Latinx) | -0.540 | 0.281 | 0.583 | 0.055 |
| | Race (other) | -0.158 | 0.255 | 0.854 | 0.536 |
| | Position (faculty) | 0.592 | 0.321 | 1.807 | 0.065 |

| | | | | | |
|---|---|---|---|---|---|
| | Position (university staff) | -0.173 | 0.340 | 0.841 | 0.610 |
| | Position (student) | 0.081 | 0.363 | 1.084 | 0.824 |
| | Region (Midwest US) | 0.368 | 0.245 | 1.444 | 0.133 |
| | Region (NE US) | 0.761 | 0.232 | 2.141 | **0.001** |
| | Region (SE US) | 0.794 | 0.289 | 2.213 | **0.006** |
| | Region (SW US) | -0.047 | 0.355 | 0.954 | 0.894 |
| | Region (West US) | 0.436 | 0.250 | 1.546 | 0.081 |
| Teaching | TGNC (yes) | -0.524 | 0.250 | 0.592 | 0.036 |
| | Race (Asian) | -0.717 | 0.246 | 0.488 | **0.004** |
| | Race (Latinx) | -0.051 | 0.324 | 0.950 | 0.874 |
| | Race (other) | -0.501 | 0.248 | 0.606 | 0.044 |
| | Position (faculty) | 1.089 | 0.692 | 2.972 | 0.115 |
| | Position (university staff) | 0.466 | 0.713 | 1.593 | 0.514 |
| | Position (student) | 0.743 | 0.732 | 2.101 | 0.310 |
| | Region (Midwest US) | 0.494 | 0.261 | 1.639 | 0.058 |
| | Region (NE US) | 0.723 | 0.235 | 2.061 | **0.002** |
| | Region (SE US) | 0.435 | 0.269 | 1.544 | 0.106 |
| | Region (SW US) | 0.411 | 0.385 | 1.509 | 0.285 |
| | Region (West US) | 0.699 | 0.271 | 2.012 | **0.010** |
| Student | TGNC (yes) | -1.283 | 0.358 | 0.277 | **0.000** |
| | Race (Asian) | -0.489 | 0.394 | 0.613 | 0.215 |
| | Race (Latinx) | 0.462 | 0.537 | 1.587 | 0.390 |
| | Race (other) | -0.601 | 0.423 | 0.548 | 0.156 |
| | Position (faculty) | -0.423 | 0.917 | 0.655 | 0.645 |
| | Position (university staff) | -0.842 | 0.945 | 0.431 | 0.373 |
| | Position (student) | -0.423 | 0.918 | 0.655 | 0.645 |
| | Region (Midwest US) | 0.228 | 0.488 | 1.256 | 0.640 |
| | Region (NE US) | 0.782 | 0.434 | 2.187 | 0.071 |
| | Region (SE US) | 2.158 | 0.681 | 8.655 | **0.002** |
| | Region (SW US) | 1.103 | 0.717 | 3.013 | 0.124 |
| | Region (West US) | 0.345 | 0.460 | 1.412 | 0.454 |

Reference groups are straight, cisgender, white, outside of academia, and outside of the US. Bold text represents statistical significance after Bonferroni corrections.

**Table S4.** Results from linear regressions for workplace inclusion and exclusion.

| Outcome | Predictor | beta | SE | p-value |
|---|---|---|---|---|
| LGBTQ+ climate inclusion | (Intercept) | 3.607 | 0.111 | **0.000** |
| | TGNC (vs cis LGBQ) | -0.211 | 0.074 | **0.005** |
| | Race (Asian) | 0.000 | 0.146 | 1.000 |
| | Race (Latinx) | -0.074 | 0.117 | 0.527 |
| | Race (other) | -0.303 | 0.122 | **0.014** |
| | Position (faculty) | 0.052 | 0.112 | 0.645 |
| | Position (university staff) | 0.036 | 0.116 | 0.754 |
| | Position (student) | -0.004 | 0.112 | 0.969 |
| | Region (Midwest US) | 0.046 | 0.110 | 0.675 |
| | Region (NE US) | 0.043 | 0.103 | 0.678 |
| | Region (SE US) | 0.005 | 0.114 | 0.967 |
| | Region (SW US) | -0.009 | 0.142 | 0.948 |
| | Region (West US) | 0.062 | 0.105 | 0.556 |
| LGBTQ+ climate exclusion | (Intercept) | 2.390 | 0.127 | **0.000** |
| | TGNC (vs cis LGBQ) | 0.162 | 0.085 | 0.058 |
| | Race (Asian) | 0.094 | 0.168 | 0.577 |
| | Race (Latinx) | 0.087 | 0.134 | 0.516 |
| | Race (other) | 0.321 | 0.140 | **0.023** |
| | Position (faculty) | -0.098 | 0.129 | 0.448 |
| | Position (university staff) | -0.044 | 0.132 | 0.740 |
| | Position (student) | 0.107 | 0.128 | 0.406 |
| | Region (Midwest US) | 0.015 | 0.126 | 0.904 |
| | Region (NE US) | -0.025 | 0.118 | 0.830 |
| | Region (SE US) | 0.009 | 0.130 | 0.944 |
| | Region (SW US) | 0.095 | 0.163 | 0.559 |
| | Region (West US) | -0.016 | 0.120 | 0.896 |

Reference groups are cisgender LGBQ, white, outside of academia, and outside of the US. Bold text represents statistical significance.

**Table S5.** Results from logistic regressions for exclusionary behavior.

| Outcome | Predictor | beta | SE | OR | p-value |
|---|---|---|---|---|---|
| Experienced exclusionary behavior at work | (Intercept) | -1.822 | 0.414 | 0.162 | **0.000** |
| | TGNC (vs cis LGBQ) | 1.319 | 0.256 | 3.738 | **0.000** |
| | Race (Asian) | -0.090 | 0.551 | 0.914 | 0.871 |
| | Race (Latinx) | -0.002 | 0.439 | 0.998 | 0.997 |
| | Race (other) | 0.679 | 0.394 | 1.971 | 0.085 |
| | Position (faculty) | -0.315 | 0.423 | 0.730 | 0.456 |
| | Position (university staff) | -0.143 | 0.423 | 0.867 | 0.735 |
| | Position (student) | 0.072 | 0.397 | 1.075 | 0.856 |
| | Region (Midwest US) | 0.338 | 0.387 | 1.402 | 0.382 |
| | Region (NE US) | -0.162 | 0.384 | 0.851 | 0.674 |
| | Region (SE US) | -0.102 | 0.440 | 0.903 | 0.816 |
| | Region (SW US) | 0.079 | 0.514 | 1.083 | 0.877 |
| | Region (West US) | -0.148 | 0.397 | 0.863 | 0.710 |
| Witnessed exclusionary behavior at work | (Intercept) | -0.895 | 0.339 | 0.409 | **0.008** |
| | TGNC (vs cis LGBQ) | 0.301 | 0.224 | 1.351 | 0.178 |
| | Race (Asian) | -0.187 | 0.487 | 0.829 | 0.701 |
| | Race (Latinx) | 0.147 | 0.352 | 1.158 | 0.676 |
| | Race (other) | 0.430 | 0.364 | 1.537 | 0.238 |
| | Position (faculty) | -0.323 | 0.345 | 0.724 | 0.349 |
| | Position (university staff) | -0.079 | 0.349 | 0.924 | 0.822 |
| | Position (student) | 0.350 | 0.332 | 1.418 | 0.293 |
| | Region (Midwest US) | 0.558 | 0.326 | 1.748 | 0.087 |
| | Region (NE US) | 0.214 | 0.318 | 1.239 | 0.501 |
| | Region (SE US) | 0.670 | 0.337 | 1.954 | **0.047** |
| | Region (SW US) | 0.370 | 0.423 | 1.448 | 0.382 |
| | Region (West US) | -0.203 | 0.332 | 0.816 | 0.540 |
| Experienced exclusionary behavior outside of work | (Intercept) | -0.487 | 0.214 | 0.614 | **0.023** |
| | TGNC (vs cis LGBQ) | 0.973 | 0.217 | 2.646 | **0.000** |
| | Race (Asian) | -0.220 | 0.448 | 0.803 | 0.624 |
| | Race (Latinx) | -0.171 | 0.349 | 0.843 | 0.624 |
| | Race (other) | -0.132 | 0.366 | 0.877 | 0.720 |
| | Region (Midwest US) | -0.079 | 0.318 | 0.924 | 0.803 |
| | Region (NE US) | -0.342 | 0.307 | 0.710 | 0.265 |
| | Region (SE US) | 0.165 | 0.331 | 1.180 | 0.617 |
| | Region (SW US) | 0.085 | 0.422 | 1.088 | 0.841 |
| | Region (West US) | -0.061 | 0.305 | 0.941 | 0.842 |

Reference groups are cisgender LGBQ, white, outside of academia, and outside of the US. Bold text represents statistical significance.

**Table S6.** Results from linear regressions for perceived discrimination.

| Predictor | beta | SE | p-value |
|---|---|---|---|
| (Intercept) | 3.587 | 0.197 | **0.000** |
| TGNC (vs cis LGBQ) | -0.879 | 0.132 | **0.000** |
| Race (Asian) | 0.313 | 0.261 | 0.230 |
| Race (Latinx) | -0.401 | 0.209 | 0.055 |
| Race (other) | -0.379 | 0.218 | 0.083 |
| Position (faculty) | 0.324 | 0.200 | 0.106 |
| Position (university staff) | 0.183 | 0.206 | 0.374 |
| Position (student) | 0.174 | 0.200 | 0.383 |
| Region (Midwest US) | -0.192 | 0.196 | 0.328 |
| Region (NE US) | -0.295 | 0.183 | 0.108 |
| Region (SE US) | -0.207 | 0.203 | 0.309 |
| Region (SW US) | -0.232 | 0.253 | 0.360 |
| Region (West US) | -0.345 | 0.186 | 0.064 |

Reference groups are cisgender LGBQ, white, outside of academia, and outside of the US. Bold text represents statistical significance.

**Table S7.** Full description, frequencies, and example quotes for each of the themes from suggestions to make workplaces and professional societies more inclusive of LGBTQ+ biologists.

| Theme | Description | % (n) | Example quote - professional society | Example quote - workplace |
|---|---|---|---|---|
| Communicate institutional support | Participant describes that their workplace or society could speak out in support of LGBTQ+ individuals (including by honoring their scientific achievements), spread awareness, address LGBTQ+ issues head-on, use inclusive and careful language, and avoid an apolitical stance. | 23.4 (126) | Participant 1444: Speak out in favor of LGBTQ+ people. Give a platform to LGBTQ+ members. | Participant 609: Take a stance against the state policies. Stop being quiet about these laws affecting healthcare for LGBTQ individuals and women. Their silence is being read as support. |
| Social support/ Mentoring | Participant describes that their workplace or society could increase social support and mentoring for LGBTQ+ individuals by having meet-up groups, affinity groups, and programming specifically for LGBTQ+ people. | 16.4 (88) | Participant 1321: Organize queer events (meet-ups, happy hours, symposia) at conferences and encourage attendance by all. | Participant 1251: The thing I would most appreciate is access to identity-specific mentorship-professional guidance from more senior LGBTQ+ researchers about how to navigate life as an out scientist. |
| Increase visibility/ representation | Participant describes that their workplace or society could increase LGBTQ+ visibility and representation through pride flags, celebrating pride, and having LGBTQ+ individuals in leadership positions. | 15.8 (85) | Participant 1567: I think promoting and celebrating LGBTQI+ visibility. I feel a real problem is simply a lack of role models because by and large LGBTQI+ people are not publicly visible. | Participant 621: Provide more visibility of LGBTQ+ research community. |

| | | | | |
|---|---|---|---|---|
| Programming | Participant describes that their workplace or society could support and train others on how to be inclusive to LGBTQ+ individuals through programming, sensitivity training, and talks by LGBTQ+ individuals. | 13.9 (75) | Participant 2016: Conferences could host workshop and conversations about how to be more inclusive in our research labs, teaching roles (especially with language!), and departmental service positions. | Participant 1321: Implement trainings about bias, gender identity, workplace conduct, and historical bias and exclusion of queer people in science. |
| Trans/nonbinary awareness | Participant describes that their workplace or society could pay attention to issues affecting transgender and genderqueer individuals, including making gender-neutral bathrooms available and ensuring the correct pronouns and names are used. | 13.4 (72) | Participant 1650: Better use of pronouns and preferred names, gender inclusive restrooms at conference, gender and surname options for forms / records. | Participant 1639: I wish that pronouns were more respected. No one really seems to care and I'm constantly misgendered. |
| Institutional action | Participant describes that their workplace or society could take institutional action to change the cisheteronormative policies including family leave, hiring processes, insurance coverage, spousal hires, and pay equity. | 7.2 (39) | Participant 1843: Inclusive award nomination processes. I can not apply for LGBTQ specific awards because that will put a target on my head. To get tenure at an R1, I need to demonstrate the that my work has an impact and awards are one potential mechanism but I don't have a strong advocate at my university who will do so. What does the award nomination and selection process | Participant 1401: Provide expansive health care benefits for LGBTQ+ employees that are well documented and easily accessible. Provide explicit leave for LGBTQ+ people when undergoing healthcare procedures. Provide financial support for fertility services, adoption, and other family-related expenses. |

| | | | | |
|---|---|---|---|---|
| | | | look like? Can you self nominate? Is the selection process sensitive to differential resources or challenges? | |
| Accountability for individuals | Participant describes that their workplace or society could curb discrimination and homophobia by improving reporting mechanisms and having consequences for harassment. | 5.6 (30) | Participant 1590: Zero tolerance policies. I'm tired of people with harmful views being allowed to continue working or not receive serious repercussions for their words or actions. It makes me feel not seen and invalidates my experiences. | Participant 1212: Sanctions need to be applied when individuals misbehave due to someone's identity/orientation. There have been reports of homophobia/transphobia from faculty members in my department, that were never acknowledged or acted upon. LGBTQIA+ people keep relying on a whisper network to help guide students towards accepting mentors, which may not be available for newly out/questioning individuals. |

| | | | | |
|---|---|---|---|---|
| Financial support | Participant describes that their workplace or society could provide financial support for LGBTQ+ individuals via grants, donations, funding, fellowships, or scholarships. | 5.4 (29) | Participant 618: LGBTQ+ students need funding and resources to find success in their respective societies. We need funding to attend events and meetings. We need the societies to incentivize participation in society events with travel support and scholarships. | Participant 1314: Offer grants, fellowships and funding schemes that acknowledge the barriers in sustaining an academic career as a LGBTQI+ individual, and work towards retaining them within the workplace. |
| Consider regional politics | Participant describes that their workplace or society could avoid hosting events in places with anti-LGBTQ+ legislation or could consider local politics when making the decision of where to hold events. | 3.9 (21) | Participant 1225: Consider the locations of annual meetings/conferences and the impact of state and local policies on LGBTQIA+ attendees and their ability to attend a feel safe. | *n/a* |
| Intersectional DEI measures | Participant describes that their workplace or society could engage in diversity, equity, and inclusion initiatives broadly (not LGBTQ-specific). | 3.9 (21) | Participant 1299: Continue to put all EDI [equity, diversity, and inclusion] issues in the forefront, be vocal about allyship and support. | Participant 407: Put more effort towards diversity and inclusion rather than leaving it up to a committee which faces lots of restrictions, and make sure that LGBTQ+ people are a part of the conversation on how this happens. |

| | | | | |
|---|---|---|---|---|
| Collect data | Participant describes that the NSF or other institutions (including their workplace or society) could collect data about LGBTQ+ identities. | 1.9 (10) | Participant 1251: I think it is critical to continue to do these types of surveys, and to collect overall demographic information about LGBTQ+ scientists in these societies. We are so often overlooked in surveys and censuses, and this makes it very difficult to quantify the challenges we face and to design strategies to address them. | Participant 1306: Collect data and work to understand how LGBTQ faculty, staff, and students are doing in programs and work. |
| Diversity | Participant describes that their society or workplace could have more diversity, including hiring a more diverse group. | 1.1 (6) | Participant 586: Have more diverse representation. | Participant 1374: Have quite a diversity of hires so we don't feel we were simply hired to improve federal workplace statistics. I'm a human being with real and complex experiences…. not just another number. |
| Confidentiality | Participant describes that their society or workplace could take care to ensure confidentiality when handling personal data, communicate why it is being collected, and not disclose the LGBTQ+ identities of members to others. | 0.7 (4) | Participant 312: [My professional society] revealed people's LGBTQ identity on [a] form to all other authors, which inadvertently outed people. More attention needs to be placed on the confidentiality of this identity. | Participant 1281: Don't spread that information [individuals' LGBTQ+ identity] to others, it is the individual's business to share, no one else's. |

| | | | | |
|---|---|---|---|---|
| Nothing | Participant describes that there is nothing that their workplace or society could do to be more inclusive, either because it is already inclusive or because it lacks the power/authority to do so. | 13.4 (72) | Participant 1791: They are doing well. There is a diversity committee with an LGBTQIAA section that is quite active and lots of working groups. | Participant 200: I don't think my workplace, in this very red state, can do much since people are entitled to their opinions. |
| Participant is unsure | Participant describes that they do not know what their workplace or society could do to be more inclusive of LGBTQ+ individuals. | 1.9 (10) | Participant 2015: I'm honestly not sure. It's a scary time everywhere and I don't know what would help right now. | Participant 1278: I am unsure. |

**Data S1. (separate file)**
De-identified dataset for analyses of all participants.

**Data S2. (separate file)**
De-identified dataset for analyses of LGBTQ+ participants.
